# Musical Aesthetic Experience Reveals Differential Effects of Professional Vocal Training on Female College Students’ Brain Function: An fMRI Study

**DOI:** 10.64898/2026.10.02.756189

**Authors:** Huimei Li, Yingxue Gao, Yao Ma, Baicai Sun, Yuwei Liu, Ruizhi Zhou, An Ren, Tifei Yuan, Xiaoqi Huang, Xia Zhang

## Abstract

Musical aesthetic experience can reveal students’ cognitive, aesthetic, and emotional responses to music. However, how professional vocal training alters brain functions in female college students with varying levels of musicianship to promote these is unknown. Here we applied fMRI, psychological measures and Goldsmiths musical sophistication index (Gold-MSI) assessments to study 27 non-music major female college students consisting of instrument group (normal pitch accuracy with instrumental training), general group (normal pitch accuracy without instrumental training) and poor-pitch group (poor pitch accuracy and without instrumental training). After 1 year of one-on-one instructional practice, we found that professional vocal training enhanced information processing related to musical cognition and pleasure during musical aesthetic experiences across all three groups, involving improvements in motor control of vocalization, auditory feedback processing, musical memory and attention together with musical pleasure. Specifically, instrument group showed activation of reward circuits, general group exhibited activation of laryngeal motor control circuits, and poor-pitch group displayed activation of sensory/sensorimotor/motor control circuits. Both pre- and post-training, the instrument group exhibited higher connectivity than the other two groups in the left amygdala–dorsal anterior cingulate cortex (dACC) and the right posterior superior temporal gyrus (pSTG)–left insula pathway. After professional vocal training, the general group showed higher connectivity in the left amygdala–dACC, whereas the poor-pitch group demonstrated increased connectivity in the right pSTG–left insula and dACC-nucleus accumbens pathways. Surprisingly, professional vocal training activated the bed nucleus of the stria terminalis (BNST), a subcortical emotion center, and connectivity with the medial prefrontal cortex associated with musical information processing. Thus, vocal training can significantly improve musical cognition and aesthetic pleasure, and facilitates neural modulation in the brain’s emotional centers.

## Introduction

As the American music educator Bennett Reimer summarized, the highest value of music education aligns with the highest value of aesthetic education, which is to improve the quality of life by enhancing people’s insight into the essence of emotions [1]. The aesthetic value of music is realized through aesthetic experiences. Aesthetic experience involves the perception of music’s intrinsic qualities and the evaluation of its beauty and effectiveness, giving rise to aesthetic judgments, aesthetic emotions and preferences being shaped by cognitive, liking and affective processes, reflecting the experiential outcome of holistic decision [2]. It involves both rational and emotional aspects, and can produce effects such as "aesthetic pleasure" or "enjoyment for its own sake" [3].

The emotions evoked by musical aesthetic experiences are unique, arising from the inherent qualities of the aesthetic object itself, and can produce feelings such as enjoyment, awe, and nostalgia [2]. The neural correlates of pleasure and displeasure induced by music exhibit overlap with the neural correlates of the emotions "sadness" and "happiness" within music, and may influence the emotional responses evoked by the aesthetic experience [4,5]. The pleasurable experience of musical aesthetic experience is accompanied by changes in autonomic nervous system responses, triggering physiological responses such as "arousal" or "chills" [6].The pleasurable enjoyment derived from music listening is closely associated with activities in the nucleus accumbens (NAc), the central hub for reward and music-related activities, together with the caudate nucleus, amygdala, orbitofrontal cortex (OFC), anterior cingulate cortex (ACC) and insular cortex, forming key components of the reward system activated by music enjoyment [7]. Recent research has found that during music listening, the bed nucleus of the stria terminalis (BNST)—a central hub in the emotional brain network—becomes activated in response to pleasurable sensations [8].

Musical skill training is another key component of school music education, it reveals the adaptive changes in the brain are associated with the practiced musical action and its behavioral outcomes, and it influences the outcomes of musical aesthetic experience. Research has shown that the extensive practice and performance of fine motor, sensory, and multimodal integration skills required for mastering a musical instrument significantly activate brain regions involved in sensorimotor and auditory discrimination during musical melodic information processing [9], particularly when formal training begins at age seven and continues for five or more consecutive years [10]. In school music education, compared to instrumental training, vocal training is more conducive to group participation, widespread adoption, and implementation. It is also more accessible for emotional release, social connection, and cognitive engagement [11].

Singing relies on the coordination of sensory and motor systems by activating brain regions involved in auditory processing, cognitive functions and memory [12]. Vocal music singing involves the respiratory system, larynx and vocal apparatus, which are coordinated by the laryngeal, diaphragm, abdominal/chest muscles and vocal muscles (e.g., lip, jaw and tongue muscles)[13]. Among these,the laryngeal motor cortex controls voluntary vocalization through direct connections with vocal motor neurons in the brainstem [14]. Existing research indicates that studies on the effects of vocal training on the brains of healthy individuals have primarily focused on cross-sectional comparisons between the general population and professional singers or opera singers studies [15,16]. However, longitudinal studies on the effects of vocal training on brain function in healthy individuals remain extremely scarce, particularly among university students aged 18 and above who have completed the post-pubescent voice change period. Previous studies have demonstrated that singing enhances musical cognition primarily through increased neural recruitment in the inferior frontal cortex (i.e., inferior frontal gyrus, frontal operculum, and anterior insula)[17]. Evidence that singing activates brain regions associated with emotional processing (i.e., ACC, hippocampus, anterior temporal pole, and bilateral amygdala) has been found only in opera singers [18]. Therefore, whether the effects of vocal training on the activation and functional connectivity of emotion-related brain regions differ across varying levels of musicianship remains an important question that requires further research.

In addition, most untrained individuals can maintain pitch accuracy within 50 cents when singing and can match the target pitch [19]. Approximately 10% to 15% of the population are ’’poor-pitch singing’’. When playing adjacent notes on a keyboard (intervals of less than a semitone), their vocal pitch deviates from the correct pitch by as much as 100 decibels or more [20]. Research has shown that vocal training can correct minor deviations in sound output and activate the experience-dependent network for auditory-motor control of singing (i.e., the superior temporal gyrus, parietal lobe, insula, and ACC) [21]. However, it remains unclear whether improvements in pitch accuracy achieved through vocal training by individuals with poor pitch accuracy lead to changes in the brain regions associated with emotion and aesthetics, as well as their functional connectivity.

In summary, during musical aesthetic experience, the brain engages in complex interactions when processing information related to musical structure and form (cognitive), aesthetics, and emotional responses [22].However, the extent to which vocal training alters brain functions in non-music-majors college students with varying levels of musicianship to promote music cognition, aesthetics and emotion is entirely unknown. Males and females possess distinct brain processing mechanisms in music perception and musical ability [23], and there are also certain differences in vocal training methods. Therefore, this study employs functional magnetic resonance imaging (fMRI), the Gold-MSI assessments [24], and psychological assessments-including the Self-Rating Depression Scale (SDS)[25], the Self-Rating Anxiety Scale (SAS)[26], the Patient Health Questionnaire-9 (PHQ-9)[27], the Pittsburgh Sleep Quality Index (PSQI)[28], and the Big Five Personality Inventory [29]-to investigate differences in whole-brain regional activation and functional connectivity during the aesthetic processing of songs (cognitive, aesthetic, emotional) among participants in the instrumental, the general group, and the poor-pitch group, both before and after one year of professional vocal training. We hypothesized that: (1) After professional vocal training, all three groups of participants will show varying degrees of improvement in musical cognition and aesthetic pleasure compared to their pre-training levels. Specifically, they will exhibit varying degrees of activation and enhancement in auditory discrimination, sensory integration, musical memory, and reward-related brain regions. (2) After the training, the three groups of participants may exhibit different changes in the functional connectivity between music-related networks (the auditory network, default mode network, central executive network, and affective network) and the rest of the brain. (3) The poor-pitch group may show a significant improvement in pitch accuracy after training, and changes may occur in brain activity (e.g., NAc, amygdala, hippocampus, ACC, and insula) and functional connectivity associated with aesthetic reward.

## Materials and Methods

### Participants

A total of 30 healthy female college first-year students (aged 18–19) from Qingdao University, all non-music majors, participated at the beginning of this study. These participants, right-handed without prior singing training and history of neurological or psychiatric disease, were divided into the instrument group (more than five years musical instrument training since the age before seven and normal pitch accuracy), general group (normal pitch accuracy without instrumental training) and poor-pitch group (poor-pitch accuracy and without instrumental training) of 10 each. To ensure the quality of instruction, teachers provide one-on-one guidance.This study was approved by the Ethics Committee Medicine College of Qingdao University. Informed consent was obtained from all participants in this study.

### Professional vocal training

The author, H.L., is a professional vocal instructor and singer who led a one-year teaching program consisting of two classes per week, for a total of 40 sessions and 4,800 minutes.The first part of the course focused on vocal training, covering breathing exercises, vocal fold closure, articulation and enunciation, as well as resonance training in the oral and nasopharyngeal cavities.The second part centered on song performance, during which participants learned and performed seven lyrical Chinese art songs, offering artistic interpretations of these works. After class, the instructor divided all participants into five groups of six, with two members each from the instrumental group, the general group, and the poor-pitch group. Each group was required to practice together once or twice a week. All participants practiced for 90 to 130 minutes each week after class, either with piano accompaniment or by humming along to audio tracks.

### Pitch-matching test

In this study, vocal teachers had each student vocally match single notes and melodic intervals (natural and altered) presented on a piano. Next, each participant recorded a song of their choice using the same microphone under identical conditions. From each unprocessed audio file, the first 50 notes were selected. Using the trial version of Melodyne 5 software, pitch accuracy was assessed with the algorithm set to "Melodic", and the Cent (CT) value was calculated for each note. The CT values ranged from F3 to B3 and from C4 to B4. Based on the results of these two tests, participants were divided into the instrumental group, the general group, and the poor-pitch group.

### MRI data acquisition

All participants underwent MRI scans both before and after the singing training at the Affiliated Hospital of Qingdao University using the Siemens Prisma 3.0T MR imaging system with a 64-channel phased-array head coil. Participants were instructed to keep their eyes closed without falling asleep and listened to the same, unfamiliar, lyrical song (Chinese) during the scanning. T1-weighted images were acquired using an MPRAGE sequence with: TR/TE = 2530/2.65 ms, TI = 1100 ms, flip angle = 7°, slice thickness = 1 mm, 224 sagittal slices, FOV = 256 × 256 mm², matrix = 256 × 256, voxel size = 1 × 1 × 1 mmt.Functional images were acquired using a GRE-EPI sequence with: TR/TE = 2000/30 ms, flip angle = 90°, slice thickness = 3.5 mm, 33 slices, no interslice gap, FOV = 224 × 224 mm², matrix = 64 × 64, voxel size = 3.5 × 3.5 × 3.5 mmt. Each fMRI run lasted 8 min and comprised 240 volumes.

### Imaging preprocessing

The preprocessing of fMRI data was conducted using the Data Processing & Analysis for Brain Imaging (DPABI, version 4.2)[30] software. The preprocessing procedures were as follows: (i) discarding the first 10 volumes; (ii) slice timing correction and realignment of images were performed; (iii) regressing out Friston-24 motion parameters (6 parameters, their previous time point, and 12 squared terms), linear drift, white matter, cerebrospinal fluid, and global signals to reduce motion and physiological noise; (iv) the individual functional images were normalized to the standard Montreal Neurological Institute (MNI) space by using Diffeomorphic Anatomical Registration Through Exponentiated Lie Algebra (DARTEL) and resampled to 3 × 3 × 3 mmt; (v) spatial smoothing using a 6 mm full width at half maximum (FWHM); (vi) temporal band-pass filtering (0.01–0.08 Hz) to remove low-frequency drift and high-frequency noise. No participant was excluded based on mean FD > 0.25 mm or more than 30% of volumes with FD > 0.5 mm.

### Regional brain activity analysis

The amplitude of low frequency fluctuations (ALFF) was used to measure regional brain activity. For each voxel, the time series was transformed to the frequency domain via FFT, and the square root of the power spectrum averaged over 0.01–0.08 Hz was computed as the ALFF value, reflecting the amplitude of spontaneous BOLD oscillations and thus local neuronal activity [31]. To enable between-subject comparisons, each voxel’s ALFF value was divided by the whole-brain mean ALFF, yielding normalized ALFF maps.

### Seed-based functional connectivity analysis

Seed-based functional connectivity (FC) was used to examine connectivity between key regions of four music-related networks—auditory, default mode (DMN), central executive (CEN), and affective (limbic and reward circuits) networks—and the rest of the brain. For the auditory network, we selected the bilateral anterior superior temporal gyrus (aSTG), posterior STG (pSTG), Heschl’s gyrus (HG), planum polare (PP), and planum temporale (PT) as seed regions defined by the Harvard-Oxford Cortical atlas. For the affective network,we selected the bilateral amygdala and hippocampus (Anatomical Automatic Labeling atlas), bilateral NAc (Harvard-Oxford Cortical atlas) and bilateral BNST [32]. For higher-order cortical regions, we used spherical seeds (6-mm radius) centered on coordinates established in prior literature to capture their functional cores and account for inter-individual variability in functional localization: the posterior cingulate cortex (PCC) [−8, −56, 26] and medial prefrontal cortex (mPFC) [−6, 52, −2] in the DMN [33]; the bilateral dorsolateral prefrontal cortex (dlPFC), left dlPFC [−43, 23, 25], right dlPFC [43, 23, 25], bilateral posterior parietal cortex (PPC), left PPC [−38, −53, 45], right PPC [38, −53, 45] in the CEN; and for the affective network, the bilateral anterior insula [−37, 25, −4] and dACC [6, 7, 43] [34,35]. Seed locations are shown in Fig S1.

For each seed, the mean time series was extracted, and Pearson correlations with all other voxels were computed, then converted to z-values using Fisher’s Z transformation. Individual FC maps were generated for each seed.

### Statistical Analysis

#### Demographics and behavioral measures

We performed one-way ANOVA in SPSS 24.0 was used to compare age, head motion, and behavioral measures (pitch accuracy, Gold-MSI, SDS, SAS, PHQ-9, and PSQI scores) among the three groups pre- and post-training. Post-hoc pairwise comparisons using Bonferroni correction to identify specific group differences. Paired t-tests examined pre-to-post changes within each group. Significance was set at p < 0.05.

#### Baseline group comparison

We conducted an ANCOVA to examine differences in pre-training ALFF and FC maps for each seed region among the three groups using Statistical Parametric Mapping version 12 software. To control for potential confounding factors, age, sex and mean FD were included as covariates. Statistical threshold was set at p < 0.005 at the voxel level and p < 0.05 at the cluster level (FWE-corrected). For FC analyses involving 27 seeds, the seed-level threshold was Bonferroni-corrected to p < 0.05/27 = 0.0018. Significant clusters were extracted for ALFF and FC values, followed byn Bonferroni-corrected pairwise comparisons to identify specific group differences.

#### Group-by-time interaction

To assess the effects of group-by-time interactions on ALFF and FC values, we used the clusters identified by pre-training differences as masks to extract the corresponding ALFF and FC values for all participants at both pre-training and post-training time points. These values were then analyzed using repeated-measures ANOVA in SPSS 24.0 to examine within- and between-group changes over time. Simple-effects tests were conducted for post-hoc within-group pre-to-post comparisons, with significance set at p < 0.05.

#### Correlation analysis

We conducted correlation analysis to examine the relationship between ALFF or FC changes before and after singing training and changes in pitch accuracy and Gold-MSI scores, as well as changes in scores of emotion and personality scales in the three groups. For variables that followed a normal distribution, Pearson correlation was used; for those that did not, Spearman correlation was applied. For exploratory purposes, the statistical significance threshold was set at p < 0.05 without multiple comparisons correction.

## Results

### Demographics of participants

Because three participants were unable to complete the training, the final study included participants in the instrumental group (n = 9), general group (n = 8), and poor-pitch group (n = 10), for a total of 27 participants. The three groups showed no significant changes in "Big Five" personality trait scores before or after professional vocal training (Table 1).

**Table 1.**
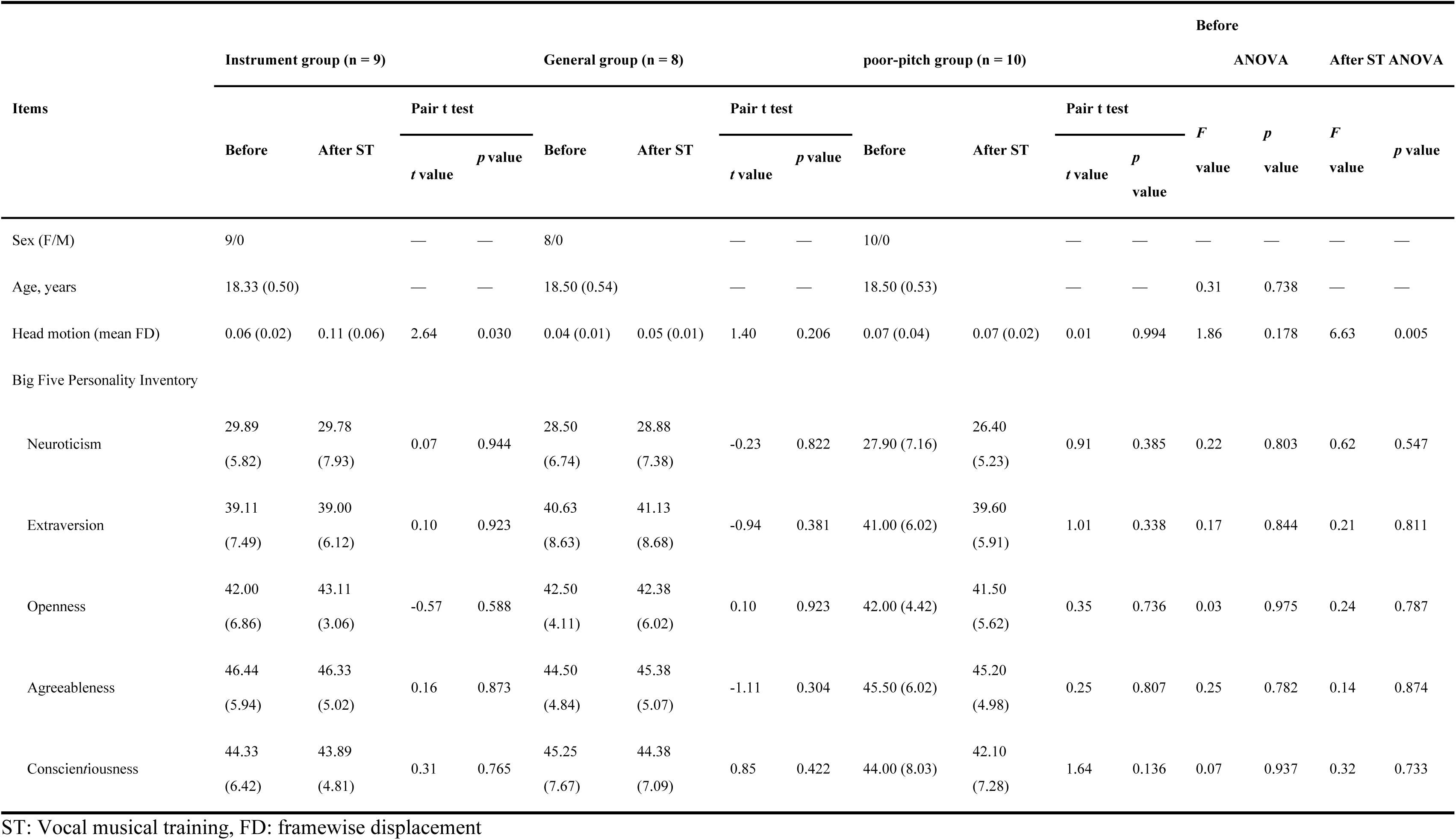
Demographics of participants.

### Pitch accuracy and training duration

All 27 participants attended at least 37 of the 40 vocal lessons. After the class, each participant practiced for an average of 115.3 minutes (standard deviation ±12.4).There were no significant differences in total training time among the three groups. Before training, there was a significant difference in pitch accuracy across the three groups. Poor-pitch group showed lower pitch accuracy than instrument and general groups. After training, the pitch accuracy of poor-pitch group significantly increased so that no significant differences were found across the three groups (Table 2).

**Table 2.**
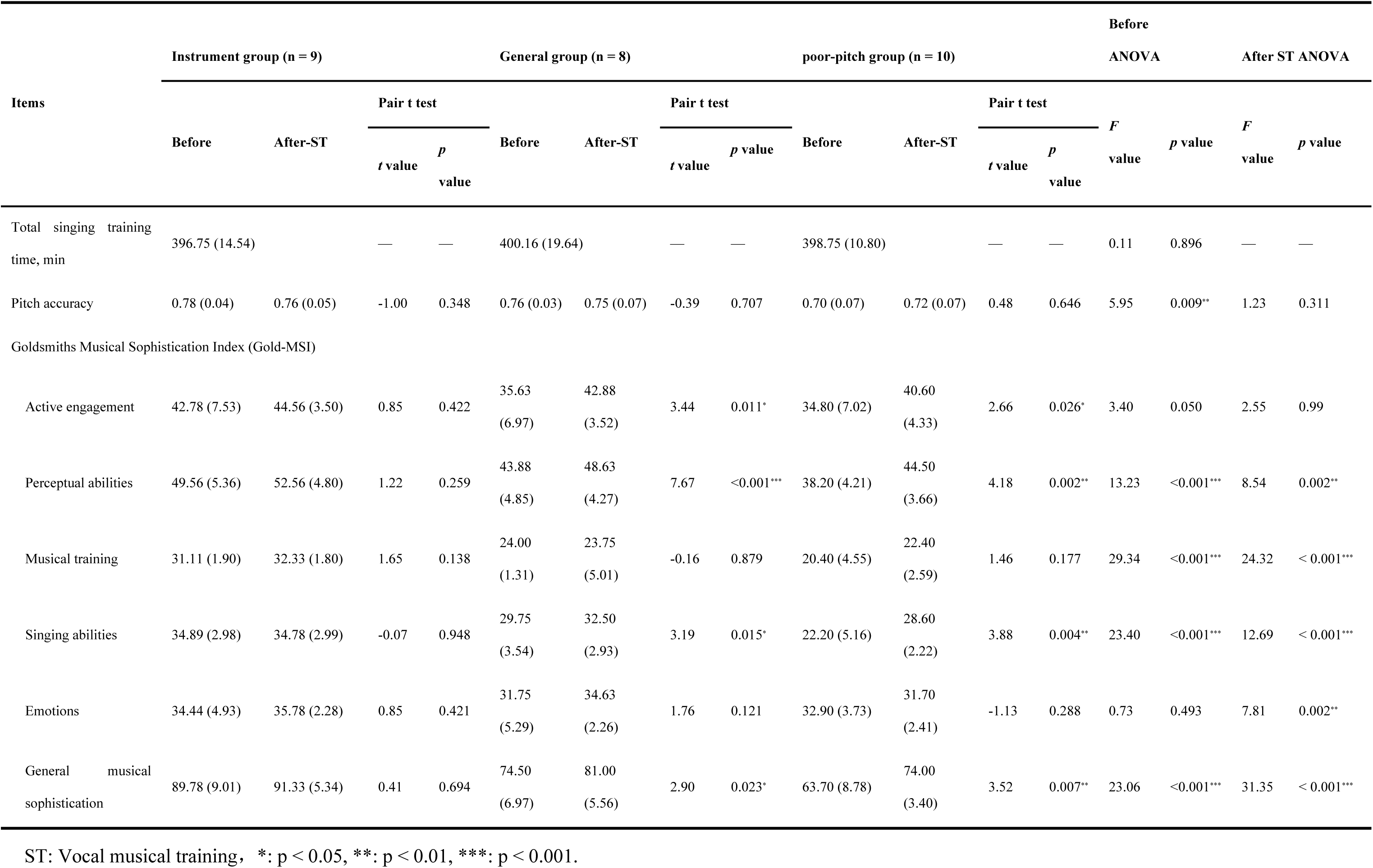
Musical measures in participants before and after professional vocal training.

| Items | Instrument group (n = 9) |  |  |  | General group (n = 8) |  |  |  | poor-pitch group (n = 10) |  |  |  | Before ANOVA |  | After ST ANOVA |  |
| --- | --- | --- | --- | --- | --- | --- | --- | --- | --- | --- | --- | --- | --- | --- | --- | --- |
|  | Before | After-ST | Pair t test |  | Before | After-ST | Pair t test |  | Before | After-ST | Pair t test |  | F value | p value | F value | p value |
|  |  |  | t value | p value |  |  | t value | p value |  |  | t value | p value |  |  |  |  |
| Total singing training time, min | 396.75 (14.54) |  | — | — | 400.16 (19.64) |  | — | — | 398.75 (10.80) |  | — | — | 0.11 | 0.896 | — | — |
| Pitch accuracy | 0.78 (0.04) | 0.76 (0.05) | -1.00 | 0.348 | 0.76 (0.03) | 0.75 (0.07) | -0.39 | 0.707 | 0.70 (0.07) | 0.72 (0.07) | 0.48 | 0.646 | 5.95 | 0.009** | 1.23 | 0.311 |
| Goldsmiths Musical Sophistication Index (Gold-MSI) |  |  |  |  |  |  |  |  |  |  |  |  |  |  |  |  |
| Active engagement | 42.78 (7.53) | 44.56 (3.50) | 0.85 | 0.422 | 35.63 (6.97) | 42.88 (3.52) | 3.44 | 0.011* | 34.80 (7.02) | 40.60 (4.33) | 2.66 | 0.026* | 3.40 | 0.050 | 2.55 | 0.99 |
| Perceptual abilities | 49.56 (5.36) | 52.56 (4.80) | 1.22 | 0.259 | 43.88 (4.85) | 48.63 (4.27) | 7.67 | <0.001*** | 38.20 (4.21) | 44.50 (3.66) | 4.18 | 0.002** | 13.23 | <0.001*** | 8.54 | 0.002** |
| Musical training | 31.11 (1.90) | 32.33 (1.80) | 1.65 | 0.138 | 24.00 (1.31) | 23.75 (5.01) | -0.16 | 0.879 | 20.40 (4.55) | 22.40 (2.59) | 1.46 | 0.177 | 29.34 | <0.001*** | 24.32 | < 0.001*** |
| Singing abilities | 34.89 (2.98) | 34.78 (2.99) | -0.07 | 0.948 | 29.75 (3.54) | 32.50 (2.93) | 3.19 | 0.015* | 22.20 (5.16) | 28.60 (2.22) | 3.88 | 0.004** | 23.40 | <0.001*** | 12.69 | < 0.001*** |
| Emotions | 34.44 (4.93) | 35.78 (2.28) | 0.85 | 0.421 | 31.75 (5.29) | 34.63 (2.26) | 1.76 | 0.121 | 32.90 (3.73) | 31.70 (2.41) | -1.13 | 0.288 | 0.73 | 0.493 | 7.81 | 0.002** |
| General musical sophistication | 89.78 (9.01) | 91.33 (5.34) | 0.41 | 0.694 | 74.50 (6.97) | 81.00 (5.56) | 2.90 | 0.023* | 63.70 (8.78) | 74.00 (3.40) | 3.52 | 0.007** | 23.06 | <0.001*** | 31.35 | < 0.001*** |

ST: Vocal musical training, \*: $p < 0.05$ , \*\*: $p < 0.01$ , \*\*\*: $p < 0.001$ .

### Musical sophistication assessment

Before professional vocal training, the instrument group showed significantly higher perceptual abilities, musical training, singing abilities and general musical sophistication scores than other two groups, while poor-pitch group showed significantly lower singing abilities and general musical sophistication scores than general group. After professional vocal training, however, both general and poor-pitch groups showed significant increases in active engagement, perceptual abilities, singing abilities, and general musical sophistication scores, whereas the instrument group did not show significant changes (Table 2).

### Psychological assessments

After professional vocal training, SAS and SDS scores in general group both decreased significantly, whereas in the poor-pitch group only the SAS scores decreased significantly. Before and after professional vocal training, there were no significant differences among the three groups in their scores on the PSQI and the PHQ-9 (Table 3).

**Table 3.** Psychological measures in participants before and after professional vocal training.

| Items | Instrument group (n = 9) |  |  |  | General group (n = 8) |  |  |  | poor-pitch group (n = 10) |  |  |  | Before |  | After | ST |
| --- | --- | --- | --- | --- | --- | --- | --- | --- | --- | --- | --- | --- | --- | --- | --- | --- |
|  |  |  |  |  |  |  |  |  |  |  |  |  | ANOVA |  |  | ANOVA |
|  | Before | After-ST | Pair t test |  | Before | After-ST | Pair t test |  | Before | After-ST | Pair t test |  | *F* value | *p* value | *F* value | *p* value |
|  |  |  | *t* value | *p* value |  |  | *t* value | *p* value |  |  | *t* value | *p* value |  |  |  |  |
| Self-Rating Depression Scale | 41.89 (8.61) | 39.22 (9.43) | -1.50 | 0.172 | 46.13 (8.94) | 40.00 (7.03) | -2.45 | 0.044* | 39.50 (10.81) | 35.40 (7.91) | -1.45 | 0.181 | 1.07 | 0.358 | 0.84 | 0.445 |
| Self-Rating Anxiety Scale | 35.89 (9.28) | 35.56 (9.11) | -0.30 | 0.776 | 40.50 (7.45) | 35.75 (7.05) | -2.49 | 0.041* | 35.60 (8.49) | 33.20 (8.56) | -2.34 | 0.044* | 0.89 | 0.423 | 0.27 | 0.764 |
| Pittsburgh Sleep Quality Index | 5.11 (3.10) | 4.89 (3.14) | -0.61 | 0.559 | 4.13 (4.64) | 3.75 (3.28) | -0.50 | 0.634 | 4.80 (2.35) | 4.30 (2.75) | -0.59 | 0.569 | 0.19 | 0.833 | 0.30 | 0.746 |
| Patient Health Questionnaire-9 | 4.67 (4.24) | 5.11 (3.72) | 0.39 | 0.705 | 3.38 (3.29) | 4.50 (3.42) | 1.20 | 0.268 | 3.10 (3.28) | 4.20 (3.39) | 2.01 | 0.075 | 0.49 | 0.620 | 0.16 | 0.850 |

ST: Vocal musical training, \*: $p < 0.05$ , \*\*: $p < 0.01$ , \*\*\*: $p < 0.001$ .

### Pre- and post-training regional activity

Before vocal training, there were no significant differences in ALFF values among the three groups. We therefore focused on within-group changes following training, using paired-sample t-tests in SPM12 (voxel p < 0.001; cluster p < 0.05, FWE-corrected). To compare training effects across groups, we further examined group-by-time interactions with a relaxed voxel threshold of p < 0.005.

After professional vocal training, specific ALFF changes in different brain regions also occurred in three groups after vocal training. Instrument group showed significantly increased ALFF in the bilateral NAc, right BNST, right globus pallidus, right thalamus, pregenual ACC (pgACC), MCC, midbrain and pons, as well as significantly decreased ALFF in vermis IX (Fig 1A, Fig S3; Table S2). General group exhibited significantly increased ALFF in the bilateral putamen, bilateral thalamus, right IFG, right ITG, left BNST, MCC and midbrain (Fig 1A, Fig S4; Table S3). Poor-pitch group displayed significantly increased ALFF in the right MTG, left cerebellum VIII, right postcentral gyrus, right IFG, right ITG, and left thalamus, as well as significantly decreased ALFF in the PCC and left cerebellum Crus I (Fig 1A, Fig S5; Table S4). Moreover, we found significant group-by-time interaction effects on the left cerebellum VIII, with poor-pitch group alone showing significantly increased ALFF in this region (Fig 1B).

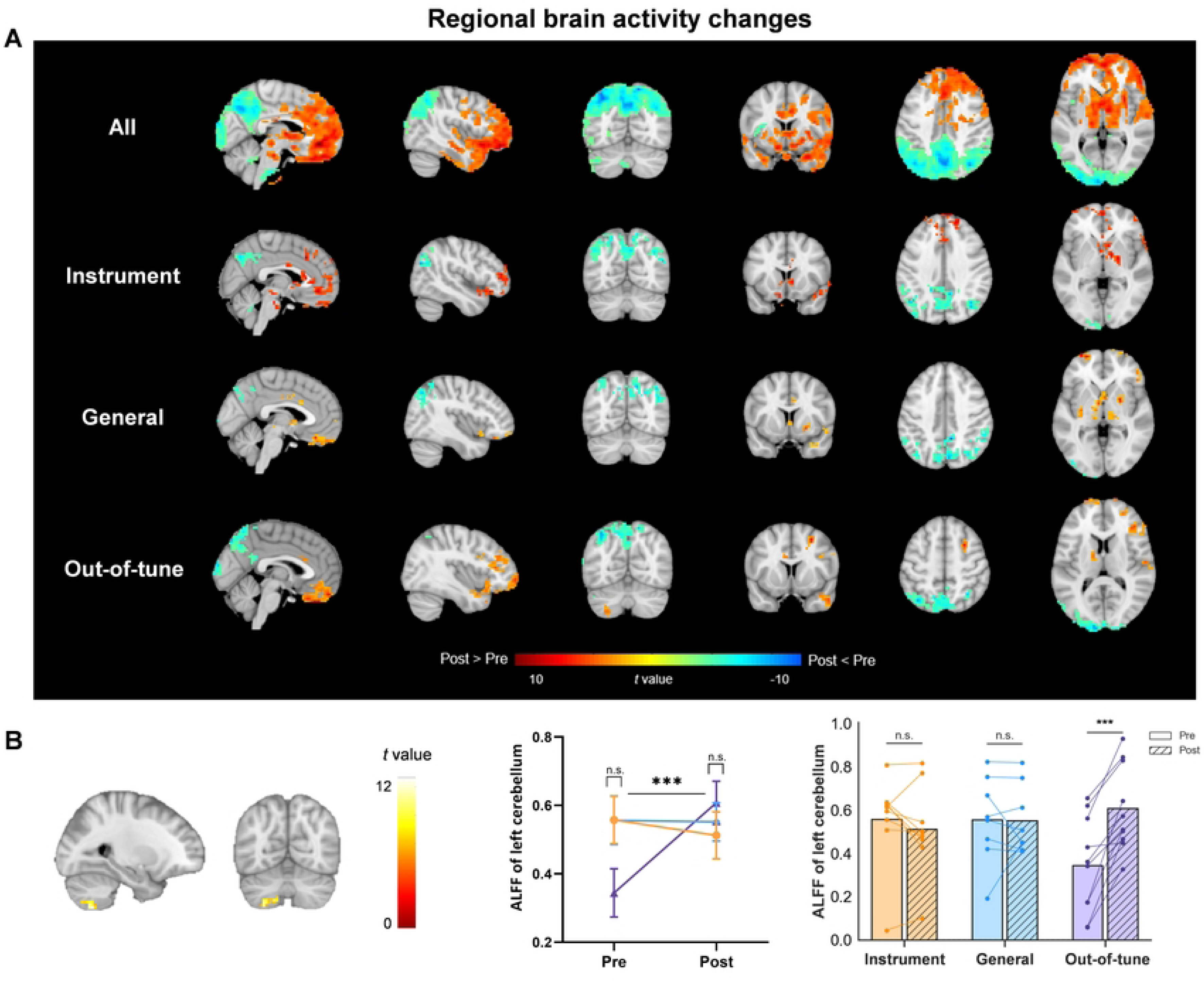

### Pre-training functional connectivity

Before vocal training, the instrument group exhibited significant higher connectivity across auditory, limbic and reward-related circuits compared to both general and poor-pitch groups (Table S5). Specifically, higher connectivity was observed in right posterior STG (pSTG)-left insula, left planum temporale-right striatum, left amygdala-dorsal ACC (dACC), and left hippocampus-left cerebellum (VI, VIIb, Crus I/II) pathways (p < 0.05, Fig 2A, Fig 3A). In comparison with poor-pitch group, instrument group showed additional connectivity alterations across reward, sensorimotor and cognitive-related regions, including higher connectivity in dACC-NAc, left BNST-supplementary motor area (SMA) and left BNST-left insula pathways, alongside lower connectivity in left anterior STG (aSTG)-right inferior parietal lobule (IPL), left NAc-left SFG, left BNST-mPFC and left BNST-OFC pathways (p < 0.05, Fig 2A, Fig 4A). Similar differences in all these pathways were also observed in general group in comparison with poor-pitch group.

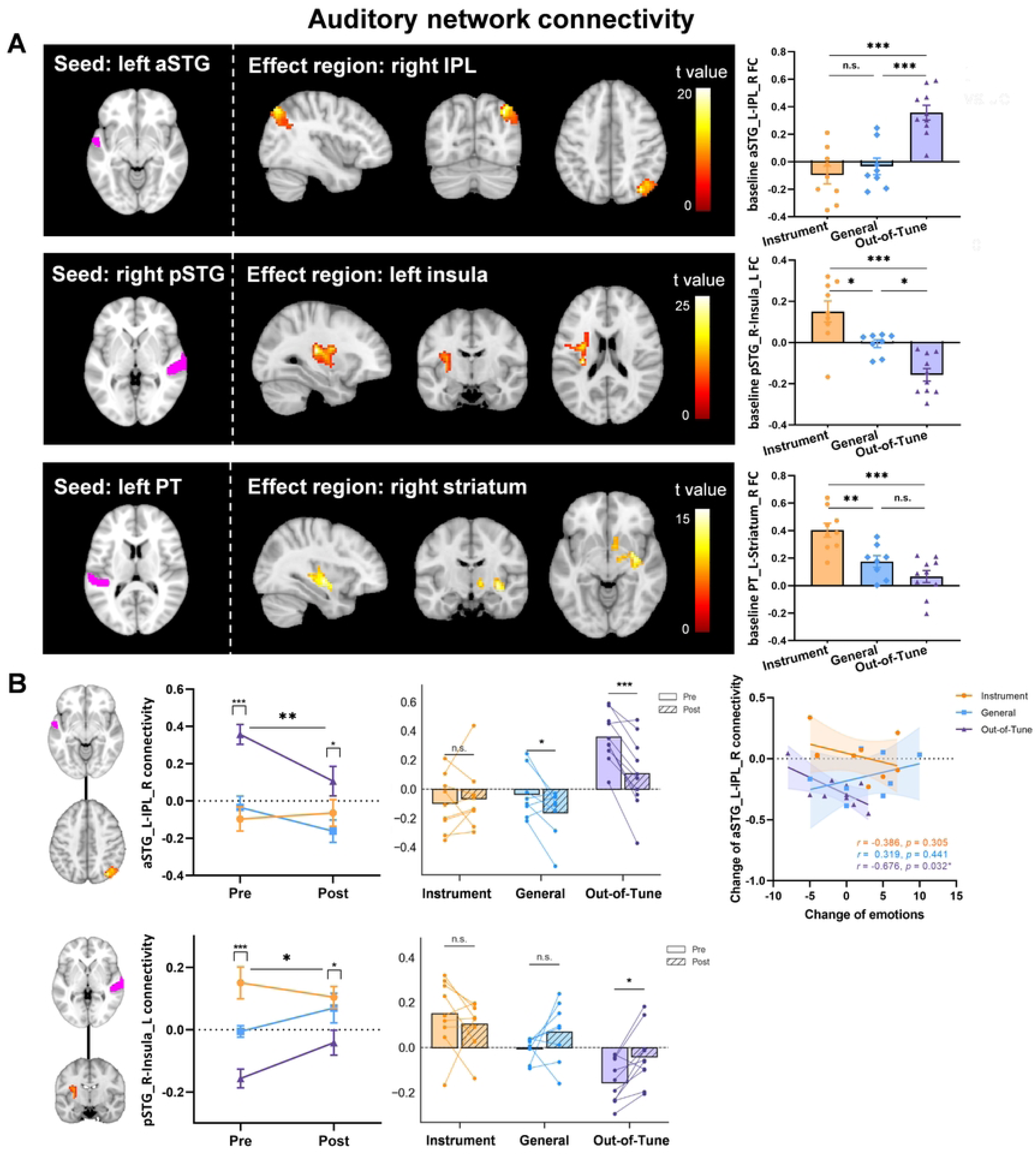

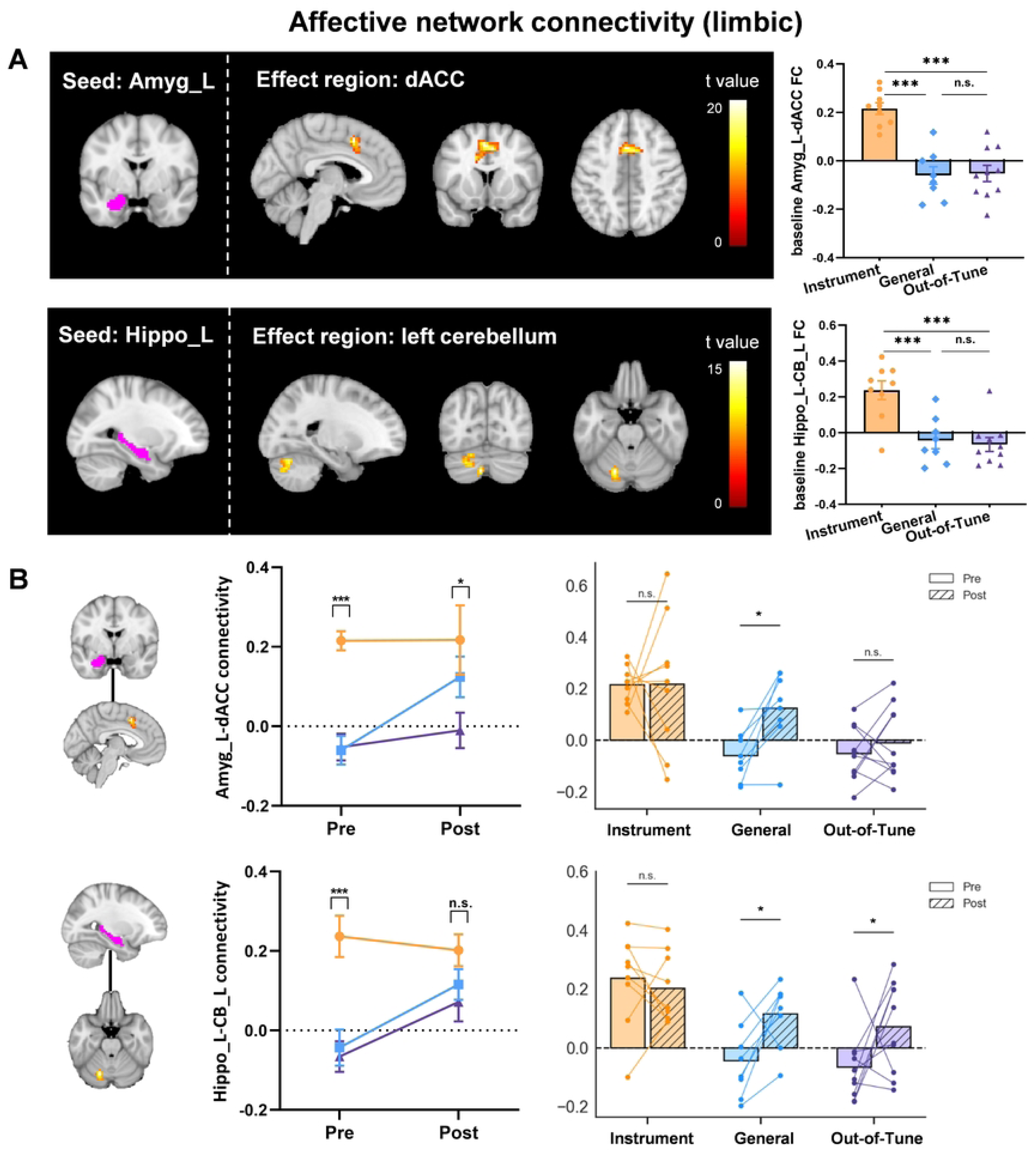

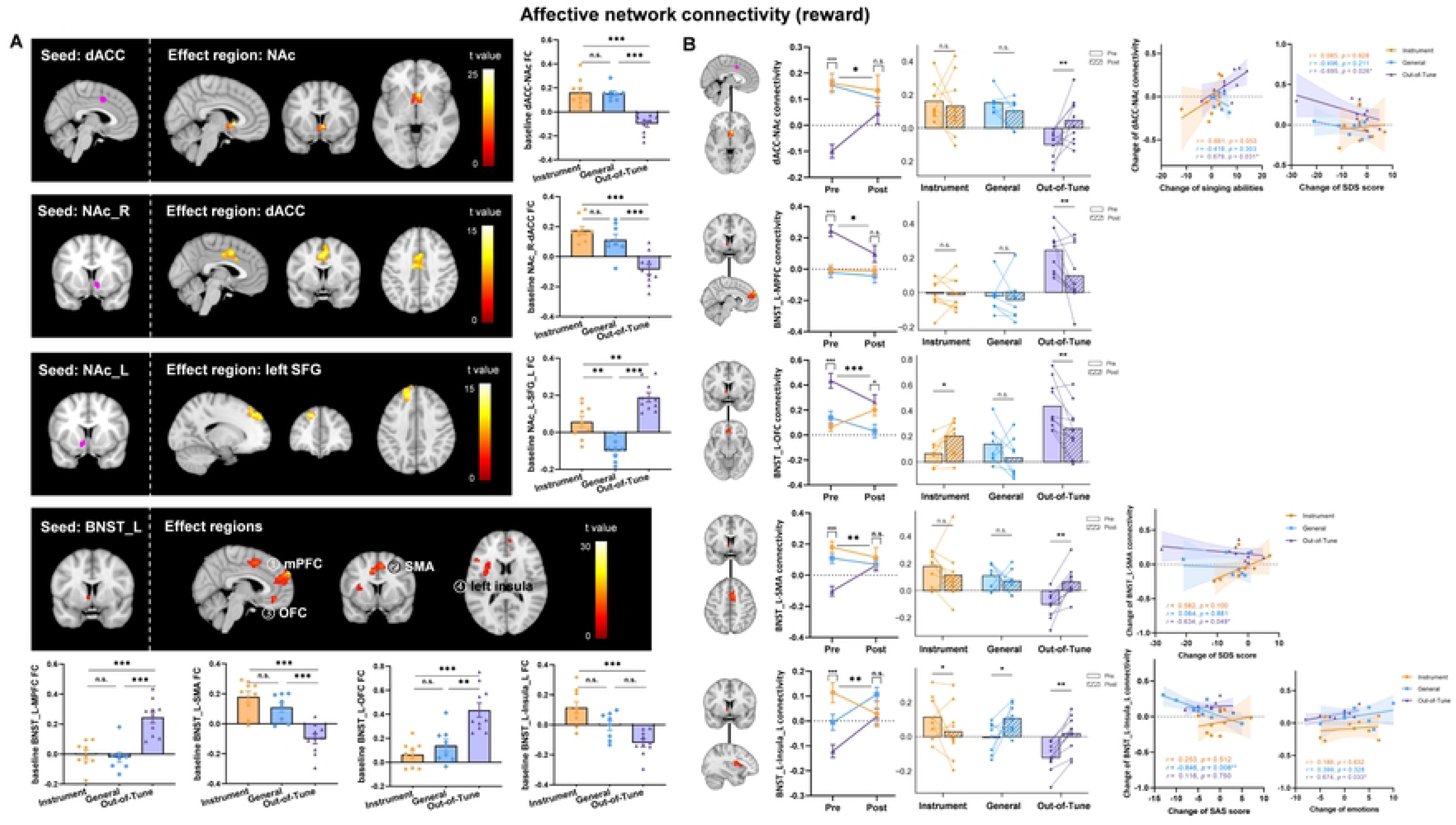

The left aSTG-right IPL, right pSTG-left insula, left amygdala-dACC, right NAc-dACC, and left BNST-mPFC connectivity remained significant after a stringent Bonferroni correction for the number of seeds selected (p < 0.05/27=0.0018). Other group differences, though did not survive this strict correction, are presented as exploratory results to provide a comprehensive overview of the training-induced changes.

### Post-training functional connectivity

After professional vocal training, we found significant group-by-time interaction effects on functional connectivity in dACC-NAcc, left aSTG-right IPL, right pSTG-left insula, left BNST-mPFC, left BNST-SMA, left BNST-OFC, and left BNST-left insula pathways (Fig 2B, Fig 4B). Although no significant group-by-time interaction effects were found for the left amygdala-dACC, left hippocampus-left cerebellum, left NAc-left SFG, or dACC-NAc connectivity, they significantly reduced in comparison with group differences observed before vocal training (Fig 3B). Post-hoc analyses revealed that, after singing training, the instrument group showed significantly increased left BNST-OFC connectivity and decreased left BNST-left insula connectivity (Fig 4B). The general group exhibited increased connectivity in left amygdala-dACC, left hippocampus-left cerebellum, and left BNST-left insula pathways, alongside reduced left aSTG-right IPL connectivity (Fig 2–Fig 4B). Post-training poor-pitch group showed increased dACC-NAc, right pSTG-left insula, left hippocampus-left cerebellum, left BNST-SMA, and left BNST-left insula connectivity, coupled with decreased left BNST connectivity with the mPFC and OFC, and left aSTG-right IPL connectivity (Fig 2–Fig 4B).

### Associations between FC and behavioral changes

We observed that increased dACC-NAc connectivity was significantly correlated with increased singing ability scores of the Gold-MSI in poor-pitch participants (Fig 4B), and that both decreased left aSTG-IPL connectivity and increased BNST-insula connectivity were significantly correlated to reduced emotion scores of Gold-MSI in poor-pitch group (Fig 2B, Fig 4B). We also found that increased dACC-NAc and BNST-SMA connections were significantly correlated with decreased SDS scores in poor-pitch group, and increased BNST-left insula connectivity was associated with decreased SAS scores in general group (Fig 4B).

## Discussion

We investigated changes in brain function before and after a year of professional vocal training in 27 healthy female college students (aged 18-19) with varying levels of musicianship but no prior vocal training. Before pre- and post-training, all participants focused on and immersed themselves in the aesthetic experience of a complete unfamiliar lyrical song during fMRI, in order to obtain fMRI data related to music information processing. We found that after training, as the three groups of participants’ musical cognition improved, they experienced greater aesthetic pleasure, and the associated functional connectivity also underwent changes to varying degrees. The poor-pitch group with pitch accuracy reached the levels of the instrumental and general groups, significantly activating brain regions associated with motor imbalance, sensorimotor processing, auditory feedback, and aesthetic perception. The general group and the instrumental group exhibited different activation patterns in the somatosensory-motor neural circuits related to vocal control. Furthermore, we were pleasantly surprised to discover that singing lyrical songs can modulate neural activity in the brain’s emotional centers. These findings confirm our research hypothesis.

This study revealed that, after professional vocal training, the three groups of female college students showed significantly enhanced activities in the following cortical regions primarily related to music cognition: the right insula, bilateral MFG, right STG, dACC and mPFC. The right insula, a key node in the brain singing network involved in auditory and somatosensory-motor feedback processing during vocalization [36], integrates multiple sensory and cognitive domains [15] and maintains and corrects pitch [21]. The MFG and right STG are respectively involved in the execution of singing, and pitch and melody processing in singing [37,38]. The dACC participates in cognitive control and makes compensatory adjustments [39]. While the mPFC activates regions related to self-referential processing and cognition during singing [40]. These cortical regions are also part of the larger“DMN”default network, which includes portions of temporal, parietal and cingulate cortices, and are crucial for adolescent introspective emotional processes [41]. Post-training participants in all three groups also exhibited reduced activities in music cognition-related brain regions, including the bilateral precuneus, bilateral IPL, left cuneus and left MOG, which are associated either with speech processing in songs [42,43] or with task-related shifts in attention [44]. These data indicate that all participants showed improved attention to music learning during vocal training.These data indicate that all participants’ attention was enhanced during the vocal training.

Each group of post-training participants also demonstrated elevated activities in specific brain regions primarily associated with music cognition. The instrumental group exhibited increased activities in the pons, right globus pallidus, and right thalamus, together with the MCC that shared elevated activities with general group. The pons connects with motor neurons controlling vocal muscles during singing and coordinates vocal muscles to produce complete vocal pattern [45,46]. The right pallidum and right thalamus are related to the cumulative degree of vocal training through professional vocal training [16,47], while the MCC reflects differences in processing musical knowledge during memory tasks [48]. The general group showed elevated activities in the bilateral putamen and bilateral thalamus, which are both associated with somatosensory motor phonation control processing during singing [10,13]. The putamen receives the strongest of all telencephalic subcortical projections from the laryngeal motor cortex [14]. The poor-pitch group displayed increased activities in the IFG, right ITG, right MTG, right postcentral gyrus, left cerebellar eighth lobe, and subcortical left thalamus, as well as decreased activities in the PCC and left superior cerebellar (Crus1). The IFG is responsible for processing basic melodies during singing and song perception [49-51], while the right ITG is associated with improvisational language and literature [52]. The right MTG is associated with pitch production during singing and music listening [53,54]. Activation of the right postcentral gyrus primarily reflects the need for non-professional singers to utilize sensory information to regulate vocal pitch control during vocalization [55]. The thalamus is involved in the motor-sensory processing of human music and language production, and shows stronger activation in professional singers during singing [56,57]. Notably, post-training poor-pitch group showed elevated activities in the left cerebellar eighth lobe, which is mainly involved in sensory-motor representation and associated with ataxia, dysmetria, dysarthria, and impaired oculomotor control [58]. In this study, the increased activity in this brain region likely alleviates articulation disorders and motor imbalance issues in singing, thereby helping to improve pitch accuracy.

In terms of functional connectivity related to music cognition, both the pre-training instrumental group and post-training poor-pitch group showed significant hyperconnectivity in the right pSTG-left insula pathway. The right STG plays a specific role in processing melody, and its stimulation leads to ’’musical stuttering’’ and pitch errors [38]. The left insula is involved in both the neural processing of vocal ecological quality and the motor coordination of articulation and phonation [59,60]. Thus, prior to vocal training, the instrumental group used sensorimotor information to regulate pitch control during vocalization, reflecting the influence of instrumental training on brain function [9]. The poor-pitch group also exhibited hyperconnectivity in this function after vocal training, which fully confirms which fully confirms that their pitch accuracy has reached normal levels and the establishment of specific functional connections related to regulating singing pitch.

Vocal training also significantly increased activities in brain regions primarily related to music aesthetics and music emotion. Thus, all three groups exhibited increased activity in the OFC, which participates in reward experiences, musical aesthetic experiences and emotional processing, mPFC, a region particularly active in aesthetic domains [18,61,62] and dACC, which primarily participates in negative emotion evaluation and exhibits heightened sensitivity to the beauty of sad music [63]. The instrumental group exhibited increased activities in the pgACC and NAc, together with shared elevations in the midbrain of general group. The pgACC coordinates autonomic and visceral motor responses and is associated with self-awareness of emotions, making it a key brain region for emotion-body response coupling [64]. The NAc, which is significantly activated during the perception of music and singing, is a primary brain region involved in the experience of musical pleasure and reward processing, and most accurately reflects an individual’s subjective feeling [6,7]. The midbrain is involved in the reward associated with singing and musical aesthetic experiences [6,40]. From this, we found that as the three groups of participants’ musical cognition improved, the brain regions associated with musical aesthetics and emotion showed varying degrees of activation.

In terms of functional connectivity related to musical aesthetics and emotion, pre-training instrumental group and post-training general group similarly exhibited increased connectivity in the left amygdala-dACC pathway. The amygdala prepares and coordinates emotional processing across other brain circuits [65,66] and is also associated with chills induced by musical aesthetic pleasure [6].The increased left amygdala-dACC connectivity suggests that vocal training helps female college students with normal pitch accuracy enhance their musical aesthetic sensitivity, leading to intense value-reward experiences.

After vocal training, both general and poor-pitch groups exhibited a significant reduction in left aSTG-right IPL connectivity. Both the left aSTG and right IPL are associated with language processing and semantic analysis during the processing of song experiences [42,67], the reduced functional connectivity likely reflects the emphasis on musicality in singing instruction, as well as the participants’ greater focus on their personal inner feelings during musical aesthetic experience, thereby enhancing their emotional and aesthetic experiences.

In addition, post-training poor-pitch group showed significantly increased dACC-NAc connectivity, whereas both pre-training instrument and general groups exhibited significantly higher functional connectivity in this pathway. The NAc and ACC are key components in enjoying music and activating the reward system [7] and enhanced connectivity in the ACC is associated with the pleasant and desirable experience of unfamiliar music [61]. Therefore, the increased dACC-NAc connectivity is likely associated with the pleasurable and perceptual enhancement of musical aesthetic experiences following improved pitch accuracy. Consistent with our findings, a recent study showed association of reduced NAc activity and connectivity with music-related anhedonia [68]. This further supports our research hypothesis that improved pitch accuracy can enhance the experience of aesthetic pleasure.

Notably, this study provides the first evidence that the emotion hub BNST and its connections showed significantly different activities before and after professional vocal training. Specifically, post-training instrument and general groups showed increased activity in the right and left BNST, respectively. The poor-pitch group pre-training exhibited excessive connectivity between the left BNST and mPFC. Post-training, this connectivity was reduced. The mPFC primarily regulates negative emotions by modulating the activity of other brain regions [63]. This reduction likely stems from the fact that vocal training improved pitch accuracy, musical cognition, and aesthetic pleasure, thereby reducing sensitivity to negative emotions. This may highlight the physiological effects of communication between the subcortical emotional processing in the BNST and cortical musical cognition functions [69]. This interaction serves to regulate emotions.

These findings reveal the mechanisms by which professional vocal training influences multiple brain functions, and provide evidence supporting the pedagogical concept of integrating music, aesthetics, and psychology within university liberal arts education.

## Author Contributions

**Conceptualization:** HL, XZ

**Methodology:** TY, XH, YG

**Investigation:** BS, YL, RZ, AR

**Visualization:** YG, YM

**Funding acquisition:** HL,XZ

**Project administration:** HL, XZ

**Supervision:** HL,XZ, XH

**Writing–original draft:** HL, YG

**Writing–review & editing:** XZ, XH,HL

